# Geography and environmental pressure are predictive of class-specific radioresistance in black fungi

**DOI:** 10.1101/2023.02.08.527471

**Authors:** Lorenzo Aureli, Claudia Coleine, Manuel Delgado-Baquerizo, Dag Ahren, Alessia Cemmi, Ilaria Di Sarcina, Silvano Onofri, Laura Selbmann

## Abstract

Black fungi are among the most resistant organisms to ionizing radiation on Earth. However, our current knowledge is based on studies on a few isolates, while the overall radioresistance limits across this microbial group and the relationship with local environmental conditions remains largely undetermined. To address this knowledge gap, we assessed the survival of 101 strains of black fungi isolated across a worldwide spatial distribution to gamma radiation doses up to 100 kGy. We found that intra and inter-specific taxonomy, UV radiation and precipitation levels primarily influence the radioresistance in black fungi. Altogether, this study provides insights into the adaptive mechanisms of black fungi to extreme environments and highlights the role of local adaptation in shaping the survival capabilities of these extreme-tolerant organisms.

**Originality statement:** Although previous studies showed the extraordinary ability of a few strains of black fungi to survive ionizing radiation, the overall radioresistance of this group of organisms has not been defined yet. Moreover, how and why radioresistance shifts across environmental gradients remain virtually unknown. Here, we collected black fungi from locations across the globe and found that biogeography shapes the responses of black fungi to environmental stress with UV light being significantly correlated with radiotolerance. Our study provides a clear picture of the boundaries of life for black fungi under ionizing radiation; further, we demonstrate, for the first time, that this ability in such microorganisms, not only is related to taxonomy, but also may be a consequence of their adaptation to various factors encountered in the environment where they live.

## Introduction

Radiation is one of the most dangerous hazards for life, among which gamma rays (i.e. high energy photons having wavelengths less than 0.1 nm) represent a serious challenge for survival and integrity of any life-forms (Dartnell, 2011). They may induce dramatic damage in biomolecules and cellular structures by direct ionization or causing radiolysis of water and, consequently, oxidative stress (Azzam et al., 2012). On Earth, there are three principal sources of gamma rays: (i) the natural radioactive nuclides, mainly ^40^K and the radionuclides from the ^232^Th and ^238^U series (Ichimiya et al., 1998; Walencik-Łata, 2022); (ii) the cosmic rays colliding with the nuclei in the atmosphere (Share et al., 2001; Wissmann et al. 2005); and (iii) human activities as in radioactive wastes, nuclear test and nuclear accident sites (Yamamoto et al., 2008; Hu et al., 2010). Albeit radioactive environments are limited only to restricted sites on our planet (Khan et al., 2020; Mousseau et al., 2020), ionizing radiation is one of the major damaging factors encountered beyond Earth (Evans et al., 2003; Dartnell et al., 2007; Roth et al., 2021); indeed, high gamma rays backgrounds are ubiquitous in interplanetary space and on the surface of most of the planetary bodies.

Among terrestrial life-forms, a small number of microorganisms have been reported to survive gamma radiation levels significantly higher than the background in the environments where they live (Gabani et al., 2013; Kwang-Woo Jung et al., 2017; Coleine et al. 2022; Coleine & Delgado-Baquerizo, 2022). Black meristematic fungi (thereafter black fungi) form an ascomycetous polyphyletic group spanning two classes (*Dothideomycetes* and *Eurotiomycetes*, mainly represented by the orders *Capnodiales* and *Chaetothyriales*, respectively) (Selbmann et al., 2020). Despite being phylogenetically distant, they share morphological characteristics as convergent adaptation to the extreme conditions where they live (Selbmann et al., 2005; Eisenman et al., 2012; Tesei, 2022). These fungi are among the most extreme-tolerant organisms on our planet, encompassing a stunning ability to survive and even flourish under prohibitive conditions, including radiation (Gorbushina et al. 2018; Selbmann et al., 2018; Coleine et al., 2022). Indeed, it has been recently reported an extraordinary resistance of these fungi to both acute and chronic exposure to a plethora of terrestrial and space relevant radiation (e.g. UV-B, X-rays, gamma rays, and cosmic rays) (Selbmann et al., 2011; Pacelli et al., 2018, 2020; Shuryak et al., 2019a; Aureli et al., 2020; Schultzhaus et al., 2020; Malo et al., 2021). These microorganisms have been investigated as possible biological means for radioprotection in contaminated environments and in manned space missions (Cordero et al., 2017; Averesch et al., 2022). Additionally, the study of radioresistant microorganisms is of astrobiological significance to unveil the survival and persistence capacity of life in highly irradiated extraterrestrial environments (Dartnell, 2011, Moissl-Eichinger et al., 2016; Horne et al., 2022).

Despite these advances, a vast bulk of questions still remains unanswered. In particular, there are three major sources of uncertainties that have precluded scientists to untangle how and why black fungi manage to resist ionizing radiation. First, most efforts have focused on a few strains (i.e. *Cryomyces, Friednmanniomyces* and *Exophiala* spp.) only, while a comprehensive study exploring the potential diverse radioresistance abilities among phylogenetically and ecologically distinct members, including those colonizing both natural and polluted anthropized niches, is still lacking. Second, most isolates come from a narrow range of environmental conditions and from local regions, while a more comprehensive study on black fungi across broader spatial distribution is lacking. Finally, very little is known on how environmental stressors, including the solar radiation levels of the regions from which these taxa are isolated, explain the actual levels of radioresistance in individual taxa.

Here, we tested the resistance of 101 black fungal strains to acute exposure to gamma radiation to relate these responses to the environmental conditions. The specimens were selected from *Dothideomycetes* and *Eurotiomycetes* classes and had a worldwide spatial distribution to cover the broadest range of different ecologies, life-styles, and geography (e.g. from Antarctica to temperate regions and from anthropic impacted to polluted sites). We aimed to: i) define the radioresistance limits of the broadest selection of black fungi tested so far; ii) identify new possible model organisms for radioprotection and astrobiology studies; iii) evaluate the potential differences in radioresistance between *Dothideomycetes* and *Eurotiomycetes*. Indeed, the first are mainly isolated from extreme dry and cold natural environments, whereas the latter show a remarkable ability to colonize hot and polluted anthropized environments (Selbmann et al., 2005; Isola et al., 2021; Tesei, 2022); iv) identify the main environmental parameters driving radioresistance in these fungi. Altogether, this information will provide new insights into the ability of black fungi to cope with high doses of ionizing radiation, paving the way for further genomic and metabolic investigations and provide a fundamental contribution to define the habitability of terrestrial environments contributing for speculations on a potential terrestrial-like life in the Solar System and beyond.

## Materials and Methods

### Strains selection

For the study, 101 strains of black meristematic fungi were selected from the Culture Collection of Fungi from Extreme Environments (CCFEE) and National Antarctic Museum Culture Collection of Fungi from Extreme Environments (MNA-CCFEE), Mycological Section of the Italian National Museum of Antarctica. The strains belong to 20 species of the order *Capnodiales* (class *Dothideomycetes*) and 7 species of the order *Chaetothyriales* (*Eurotiomycetes*) (Table S1). The strain selection was performed to study phylogenetically distinct members of meristematic black fungi and to cover a range of different environmental conditions (Table S2).

### Exposure conditions

Fungal colonies of the selected strains were incubated in triplicate on Malt Extract Agar (MEA) medium in Petri dishes according to the temperatures recorded in the localities from which the strains were isolated and the optimum growth temperature and growth rate recorded in representative strains of the tested species in previous laboratory tests: 15 °C for 3 months for *Dothideomycetes* and 25 °C for 1 week for *Eurotiomycetes* (Selbmann et al., 2005; Egidi et al., 2014). After the growth, the colonies were retrieved from each replicate and separately desiccated under laminar flow in a sterile cabinet. Dried colonies from each triplicate were exposed to gamma radiation under ambient conditions using a ^60^Co source at the Gamma Irradiation CALLIOPE Facility (ENEA Casaccia Research Centre, Rome) (Baccaro et al., 2019). The irradiation was performed at doses of 0.1, 0.5, 1, 2, 3, 5, 15, 30, 50, and 100 kGy through different exposure times at a dose rate of 1.12 kGy(air)/h. To obtain control samples, colonies from each desiccated replicate were not exposed to gamma radiation but maintained under the same environmental conditions as the irradiated samples.

### Survival assessment

The survival of microorganisms after exposure to each gamma radiation dose was measured using the colony-forming unit (CFU) test. Three colonies from each replicate exposed to gamma radiation were separately rehydrated in NaCl 0.9% solution for 72 h at 25 °C and 15 °C for *Eurotiomycetes* and *Dothideomycetes*, respectively. After rehydration, 0.1 mL of each cellular suspension at a concentration of 15,000 CFU/mL was plated in triplicate on MEA medium in Petri dishes. The dishes inoculated with *Eurotiomycetes* strains were incubated for 1 week at 25 °C, whereas dishes with *Dothideomycetes* strains for 3 months at 15 °C. The survival of microorganisms at each dose was expressed as the ratio between the mean of colonies scored in the exposed replicates and the mean of colonies scored in non-exposed samples. The decimal reduction dose (D10) value, defined as the absorbed radiation dose required to inactivate 90% of the microbial population, was estimated for each strain from its survival curve. To calculate D10 values, the best curves describing the survival of each strain were determined by fitting the exponential function (y = exp^(-bx)^) to the observed data through the Python SciPy package V.1.9.0. The curve fits were evaluated through the χ^2^ test by using the Python code.

### Intraspecific differences in radioresistance

Among the selected species, *Exophiala xenobiotica, Recurvomyces mirabilis, Meristemomyces frigidus, Elasticomyces elasticus, Extremus antarcticus, Friedmanniomyces endolithicus*, and *Cryomyces antarcticus* species were represented by more than one strain. To quantify the radioresistance variability in species including more than one strain, the coefficient of variation (CV) was calculated through the Python SciPy package. The one-way ANOVA post hoc Tukey HSD test and t-test were performed through Python bioinfokit package V.2.1.0. to assess the difference in D10 values between strains of *E. elasticus, M. frigidus, R. mirabilis* and *F. endolithicus* collected from distinct localities and among the mean D10 values calculated in the resistance groups.

### Environmental metadata acquisition

The ecological bases of the resistance to ionizing radiation was investigated by considering 27 environmental parameters obtained from the localities where the strains were isolated: Annual Mean Temperature (AMT); Mean Temperature of the Warmest Quarter (MTWAQ); Mean Temperature of the Coldest Quarter (MTCQ); Annual Precipitation (AP); Precipitation of the Wettest Month (PWEM); Precipitation of the Driest Month (PDM); Precipitation Seasonality (Coefficient of Variation) (PS); Precipitation of the Wettest Quarter (PWEQ); Precipitation of the Driest Quarter (PDQ); Precipitation of the Warmest Quarter (PWAQ); Precipitation of the Coldest Quarter (PCQ); Mean Diurnal Range (Mean of monthly (maximum temperature - minimum temperature)) (MDR); Temperature Seasonality (standard deviation ×100) (TS); Maximum Temperature of the Warmest Month (MTWAM); Minimum Temperature of Coldest Month (MTCM); Temperature Annual Range (calculated as MTWAM - MTCM) (TAR); Mean Temperature of the Wettest Quarter (MTWEQ); Mean Temperature of the Driest Quarter (MTDQ); Isothermality (calculated as MDR/TAR) (×100) (IT); solar radiation; UV index; biome type (biome); isolation source; country of isolation (country); Köppen-Geiger climate classification subgroup (KG climate); class to which the strains belong (Class); the coordinates of the localities where the samples were collected. The environmental data were available for 86 of the 101 strains tested in the study, related to 36 different localities (Supplementary Table 2, Fig. 2).

**Figure 1.**
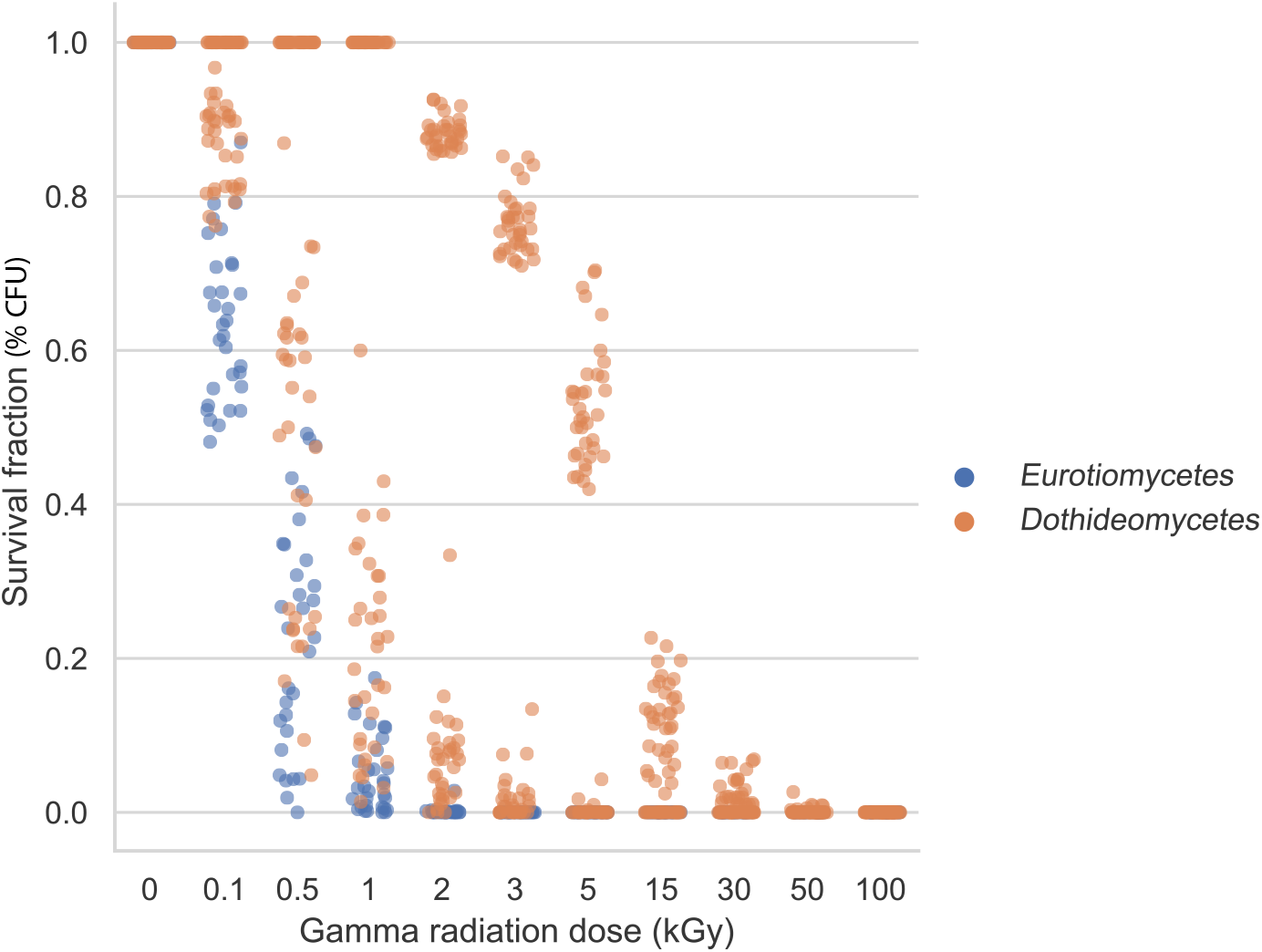

**Figure 2.**
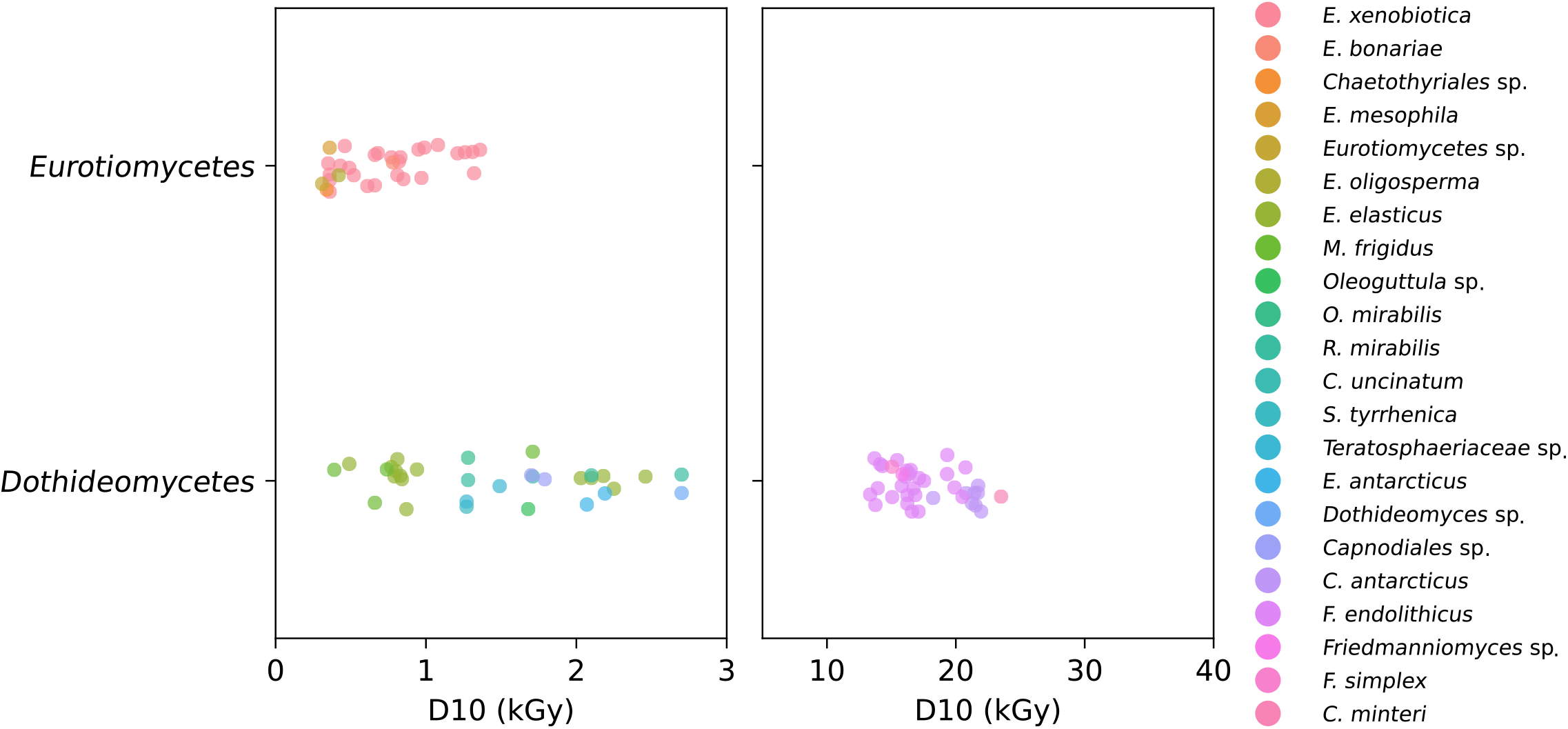

### Identification of the most important environmental parameters and statistical analyses

To identify the environmental parameters that are linked to the radioresistance of the strains, a classification predictive model was constructed through the eXtreme Gradient Boosting (XGBoost) ensemble tree-based learning algorithm. The algorithm was implemented using the Python XGBoost package V.1.6.2. Categorical data (biome, isolation source, country, KG climate, Class) were converted through ordinal encoding pre-processing. Similarly, the D10 values were grouped into 9 categories and transformed into an ordinal variable. We then estimated the importance of each environmental parameter by evaluating the mean decrease in the model score when the values of the parameter were randomly shuffled in 1000 repeats through the permutation feature importance approach by using the Python ELI5 package V.0.13.0. The XGBoost model and the permutation were first performed considering all the strains with environmental data available. Successively, the analyses on subgroups consisting of *Dothideomycetes* and *Eurotiomycetes* were performed to identify predictors of radioresistance in strains from the two classes. Finally, the 24 strains of *F. endolithicus* were considered in relation with environmental data to identify parameters that may influence intraspecific differences in radioresistance. The *F. endolithicus* species was selected due to the availability of environmental data observed in 9 distinct localities where the strains were collected. Along with *F. endolithicus, C. antarcticus, E. elasticus* and *E. xenobiotica* were the most represented species (8, 10 and 30 strains, respectively). However, the lack of environmental data related to part of *C. antarcticus* and *E. elasticus* strains did not allow the construction of a model, whereas most of *E. xenobiotica* strains were isolated from the same locality.

To examine the differences in mean decreases in the model score obtained for each environmental variable, we performed the one-way ANOVA post hoc Tukey HSD test through the Python bioinfokit package V.2.1.0. Spearman’s rank correlation coefficient and p-value were calculated through the Python SciPy packageV.1.9.0.

## Results

### Survival of fungal strains after gamma radiation exposure

The ability to generate colonies after gamma radiation exposure revealed a non-linear effect of radiation on black fungi cell survival, which was highly dependent on taxonomy. Indeed, three distinct responses can be observed from the dose-survival curves of *Eurotiomycetes* and *Dothideomycetes* strains, despite all exhibiting a similar trend that can be described by the exponential curve y = exp^(-bx)^ (Fig. S1-S3). Specifically, in all *Eurotiomycetes*, a threshold in cell survival was reported between 0.1 and 0.5 kGy, with the survival fraction reaching values below 50%. However, a drastic change in the slopes occurred beyond 0.5 kGy, thus determining a slower decrease in cell survival until reaching the maximum resistance doses of each strain (Fig. 3).

**Figure 3.**
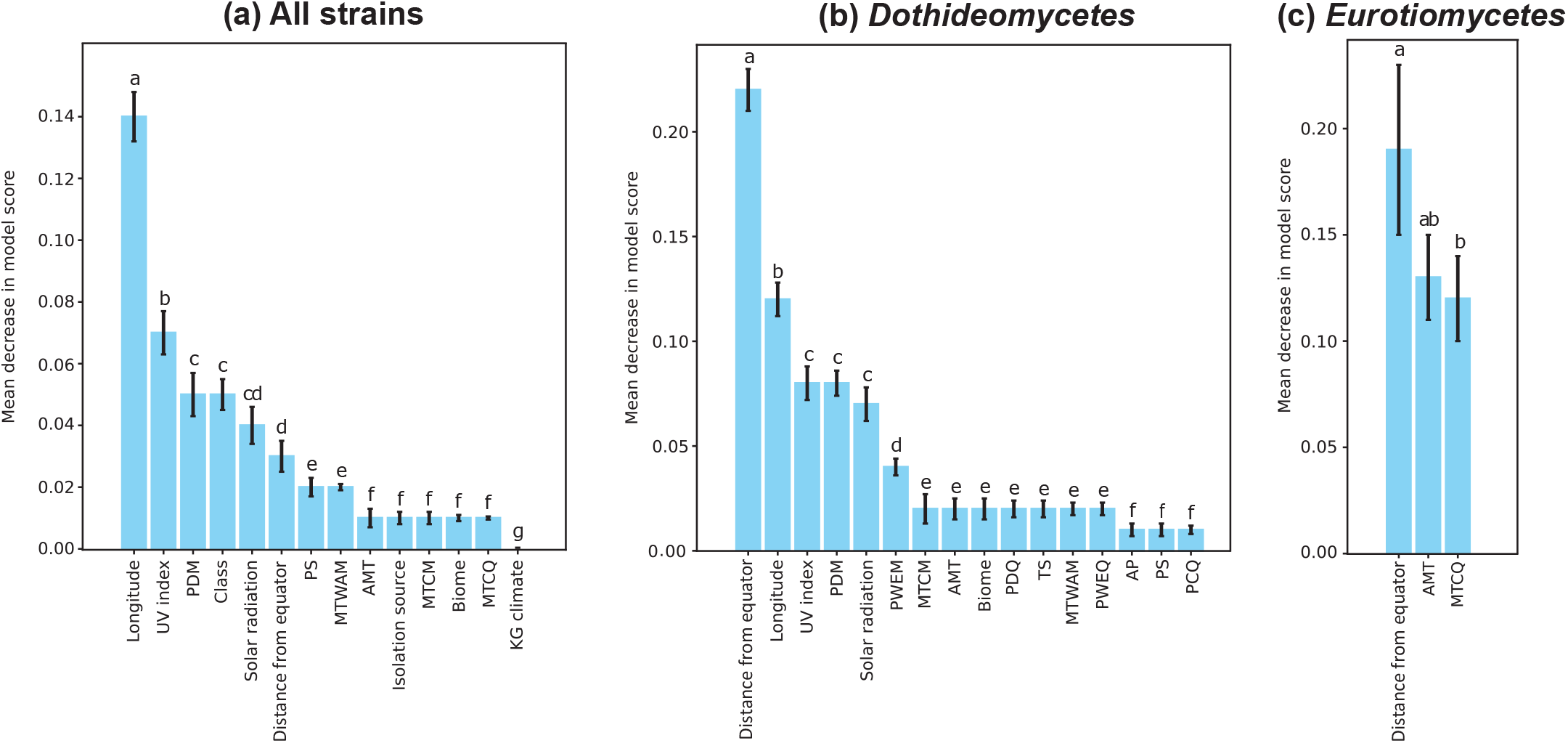

On the other hand, two different general responses to gamma radiation were shown among *Dothideomycetes* strains. The first observed response was displayed by 33 strains. Although the slopes of the curves among these strains were highly variable, they all showed a drop in cell survival from the dose of 0.5 kGy followed by a slower decrease while approaching the maximum doses of resistance. Instead, the remaining 37 strains, that included exclusively the totality of *Friedmanniomyces* and *Cryomyces* strains, showed a general drop in the cell survival between 3 and 15 kGy, followed by a slight decline in survival fraction (Fig. 3).

The D10 values obtained from the survival curves ranged from 0.30 (*Eurotiomycetes* sp. CCFEE6388) to 23.5 kGy (*C. minteri* MNA-CCFEE5187) (Table S1). Although marked differences were observed in the survival among the studied strains, colonies from most samples were able to grow after exposure at doses higher than 0.1 kGy. Furthermore, except for *E. xenobiotica* CCFEE5985, all strains formed colonies at doses higher than 1 kGy, with overall higher survival fractions observed in the *Dothideomycetes* strains (Fig. 4).

**Figure 4.**
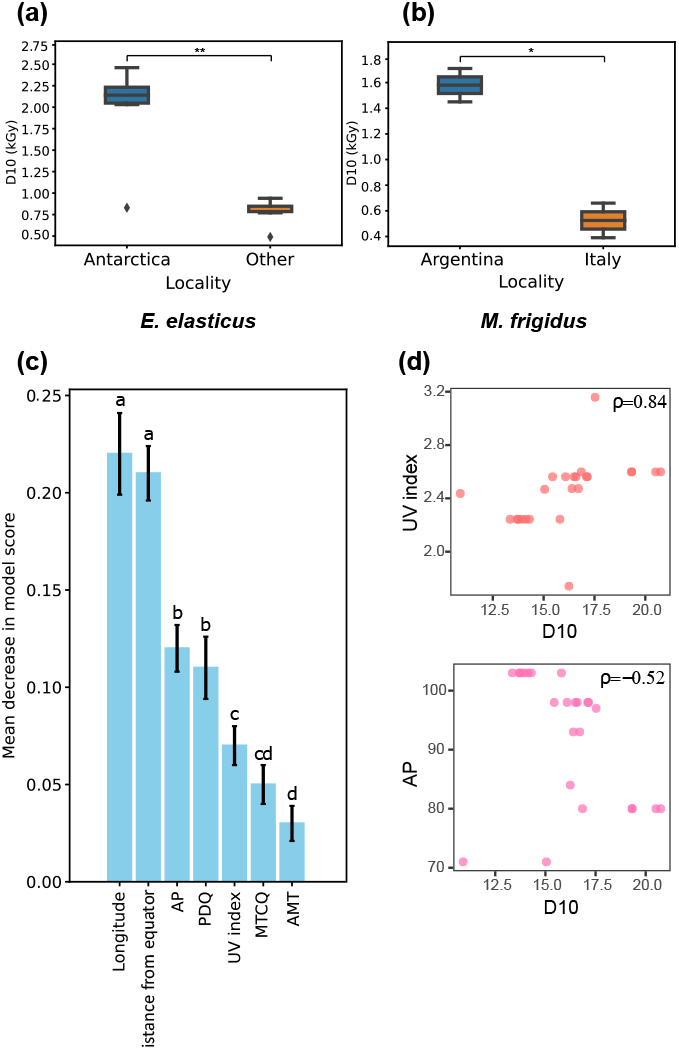

In *Eurotiomycetes*, the mean D10 was 0.8 ± 0.3 kGy with a CV 0.45. The values of the group were distributed between 0.3 and 1.4 kGy (Fig. 4), shown by *Eurotiomycetes* sp. CCFEE6388, *Chaetothyriales* sp. CCFEE6169, and *E. xenobiotica* CCFEE6180, respectively (Table S1). Moreover, only the strains *E. xenobiotica* CCFEE5819, CCFEE6180, CCFEE6182, CCFEE6196 and CCFEE6237 showed D10 above 1 kGy. The maximum resistance dose of 3 kGy was recorded for the strain *E. xenobiotica* CCFEE5877, whereas 21 out of 31 strains showed the ability to grow at the maximum dose of 1 kGy (Table S1).

In contrast, high variability in radioresistance was observed among *Dothideomycetes*, whose values ranged between 0.4 and 23.5 kGy, observed in *M. frigidus* CCFEE5401 and *C. minteri* MNA-CCFEE5187, respectively (Fig. 4). The D10 values showed a discontinuous distribution that defined two groups of radioresistance in *Dothideomycetes*. The first group was represented by strains exhibiting D10 below 4 kGy (mean D10, mean 1.5 ± SD 0.7 kGy), whose values partially overlapped the values found in *Eurotiomycetes*. However, all the strains of the group survived at doses of 2 kGy or higher, with 12 strains reaching the maximum survival dose at 5 kGy. This group consisted of all tested species of *Dothideomycetes*, with the exception of species belonging to *Cryomyces* and *Friedmanniomyces* genera. The D10 values of the second group ranged between 13.4 and 23.5 kGy, making these strains the most resistant among the tested black fungi (mean D10 17.6 ± 2.9 kGy, CV 0.15). Furthermore, all the strains of the second group formed colonies at a dose of 15 kGy or higher, with 10 strains reaching the maximum survival dose at 50 kGy (Supplementary Table S1).

### Relation between radioresistance and environmental parameters, taxonomy and geographical distribution

We further investigate the correlation between environmental and spatial conditions with the capacity of black fungi to withstand radiation. To investigate the ecological bases of the radioresistance in black fungi, we considered the importance of geography and a set of crucial environmental parameters occurring in the localities from which the strains were isolated in predicting the D10 values. The permutation feature importance scores obtained from the models indicated the spatial distribution (i.e. the longitude and the distance from the equator) of the localities as the major predictors of the radioresistance while considering the strains altogether and *Dothideomycetes* separately, although the relative importance of the two variable varies between them. Among the environmental variables, the main predictors of the radioresistance considering both the strains altogether and *Dothideomycetes* only were those related to the solar radiation exposure (i.e. UV index and solar radiation) and precipitation (i.e. precipitation in the driest month (PDQ) and precipitation seasonality (PS)) (Fig. 5a, b).

**Figure 5.**
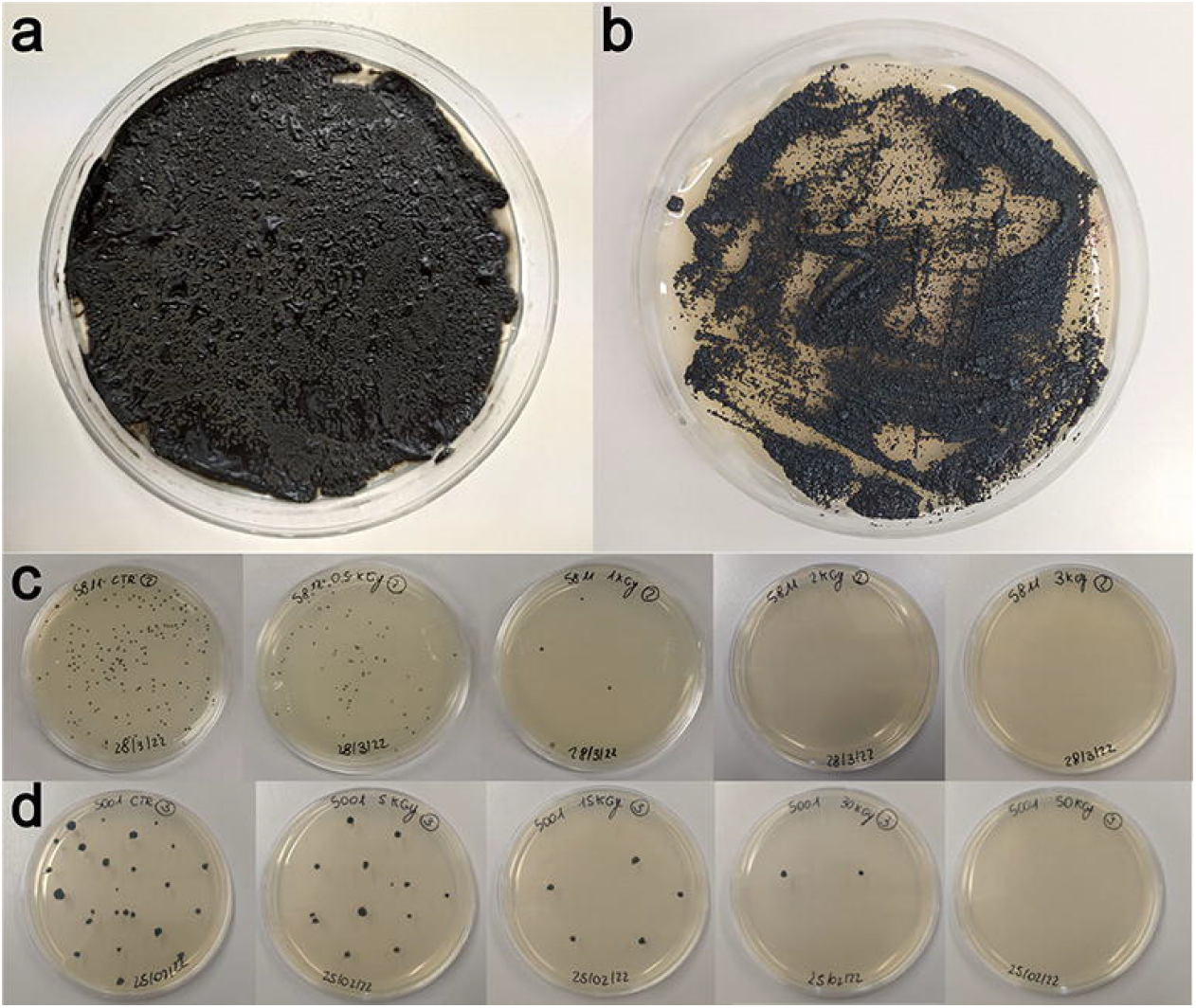

Unlike *Dothideomycetes*, strains of *Eurotiomycetes* were collected from both environments directly exposed to natural factors (e.g. surfaces of monuments) and anthropogenic and polluted environments (i.e. fuel tanks). When considering only strains exposed to natural environmental factors, parameters linked to temperature (the annual mean temperature (AMT) and the mean temperature of coldest quarter (MTCQ)) and the distance from the equator were the most important variables, whereas the solar radiation exposure and precipitation levels were shown not to play a pivotal role in radioresistance prediction for this class (Fig. 5c).

### Intraspecific differences in radioresistance and their correlation to environmental factors

In our study, 7 fungal species are represented by more than one strain (Table S1). Among them, the highest intraspecies variation in the survival data was observed in the *E. elasticus* and *M. frigidus* strains (CV 0.5 and 0.6, respectively), whereas *F. endolithicus* strains had a relatively lower CV (i.e. 0.1). In *E. elasticus*, we observed a significantly higher radioresistance (D10 above 2 kGy) in strains isolated from Antarctica compared to those isolated from temperate regions (D10 below 1 kGy) (Fig. 6a). Similar responses were observed among the strains of *M. frigidus*, whose strains from Argentinian Andes were observed to be significantly more resistant than the strains from Italy, with a difference in D10 of about 1 kGy among the two groups (Fig. 6b). These data indicated that geography and the local environmental conditions may influence the level of radioresistance in these fungi.

**Figure 6.**
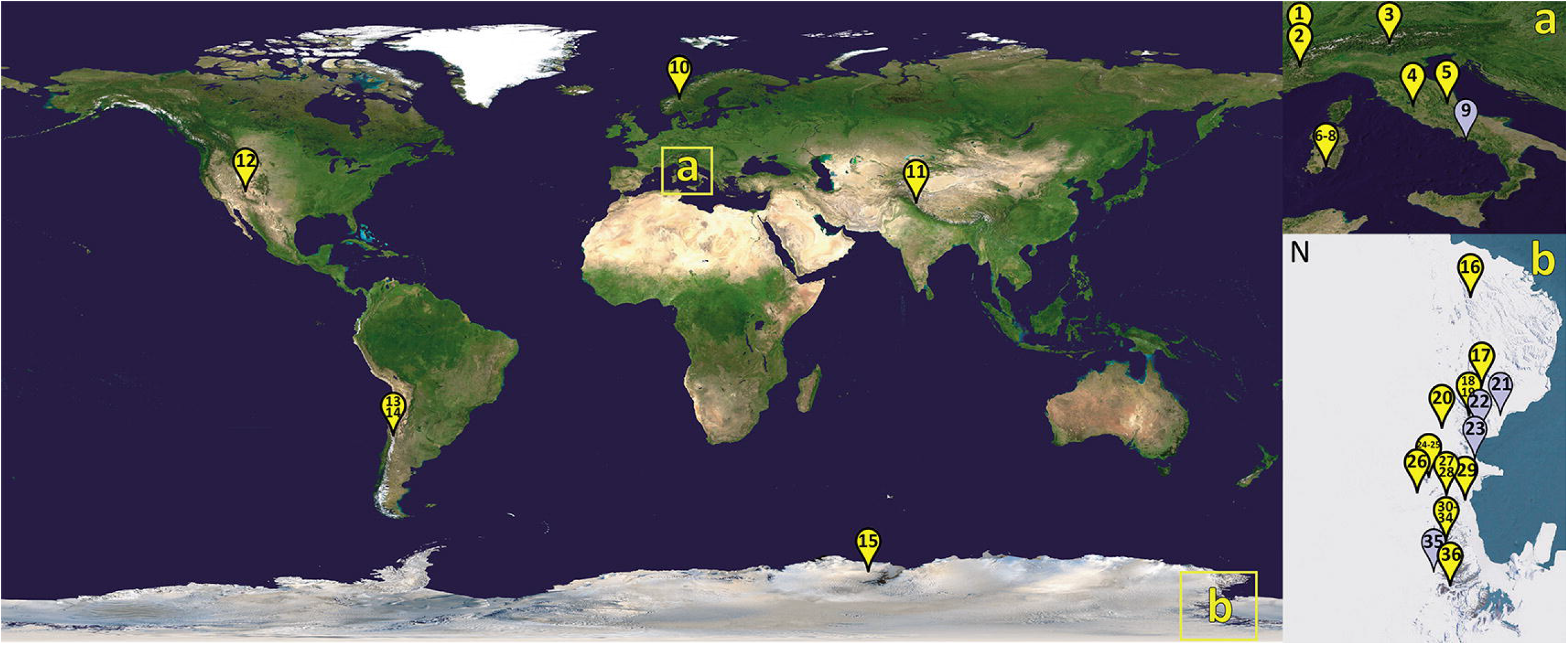

*F. endolithicus* species was represented by 27 strains isolated from 11 different localities in Antarctica (Fig. 1, Table S1). The analysis showed the geography, the precipitation parameters (AP and PDQ), the UV index, the temperature (AMT and MTCQ) as the major predictors of the radioresistance in *F. endoliticus* strains (Fig. 6c). The relationship between AP, PDQ and UV index and the radioresistance was confirmed by the Spearman’s correlation (Fig. 6d).

## Discussion

Major unknown persist about how and why different black fungi taxonomies withstand radiation across large environmental gradients worldwide. Here, the ability of 101 *Dothideomycetes* and *Eurotiomycetes* black fungi to form colonies after acute exposure to gamma radiation was assessed to define the radioresistance limits of the broadest range of these extreme-tolerant guilds tested so far. Moreover, we correlate this radioresistance with environmental and spatial conditions supporting black fungi across global environmental gradients. We showed that taxonomy is a critical component when explaining the capacity of black fungi to resist radiation. Moreover, environmental conditions were critical in explaining the variation in blackfungi radioresistance with UV light being an important predictor. These results suggest that blackfungi radioresistance levels are in consistency with the environment surrounding these organisms.

The amplitude of radioresistance capacity varied considerably among black fungi in our wide selection; many of them were able to form colonies after gamma radiation exposure doses from 0.5 to 5 kGy. Yet, some others were still viable when exposed at the extraordinary value of 50 kGy. D10 also varied considerably among strains tested, ranging from 0 kGy up to above 23 kGy, and the species falling into two distinct categories. In this study, we found a general higher aptitude to cope with ionizing radiation in dothideomycetous black fungi; in fact, the capacity to grow after lethal dose exposition over 5 kGy was an exclusive prerogative of few *Dothideomycetes*, indicating a class-tendency throughout this ability. In addition, the highest radioresistance observed (up to 50 kGy) was displayed specifically by black fungal *Dothideomycetes*, which have been isolated in the driest and coldest desert on Earth only: the ice-free areas in continental Antarctica, including the the martian analogue McMurdo Dry Valleys. Based on this evidence, we investigated in deeper detail the origin of this difference studying the effect of various environmental factors in shaping radioresistance tendency in black fungi, taking into consideration that dothideomycetous do prefer highly stressing cold-dry natural environments, while eurotiomycetous spread preferentially in hot or anthropic impacted or polluted environments.

Our study showed that biogeography, sun exposure, and the levels of precipitation were the major predictors of D10 in strains exposed to natural parameters, indicating that the ability to cope with ionizing radiation may be related to the adaptation to other environmental stressors (Shuryak et al., 2019a). Indeed, factors encountered in the extreme environments such as UV exposure, drought, and low and high temperatures can also cause oxidative stress, requiring the activation of cellular mechanisms similar to those that mediate the responses to ionizing radiation (Leprince et al., 2010; Lushchak, 2011; Kostadinova et al., 2012; Braga et al., 2015; Mejía-Barajas et al., 2017; Gostinčar et al., 2018). The burden of skills for the success in such environments, therefore, may represent a pre-adaptation for radioresistance in these fungi (Etemadifar et al., 2016; Sharma et al., 2017; Lim et al., 2019; Coleine et al., 2022). In fact, although exhibiting overall high radioresistance, none of the strains tested in our study were isolated from radioactive environments; either, the environmental background radiation should not have represented an evolutionary driver for this ability. In our study, the adaptation of black fungi to a broad range of different habitats may explain the large diversity that we observed in their radioresistance. In addition to interspecific fluctuation in radioresistance, the distinct localities may have driven the intraspecific variability in different species by inducing the local adaptation to various environmental factors (Ellison et al., 2011; Gladieux et al., 2014; Branco, 2019). Consistently with this hypothesis, the highest intraspecific variability observed was exhibited by strains of *E. elasticus* and *M. frigidus*, whose distribution spans different continents, which was highly correlated to increasing environmental pressure; indeed, the most resistant strains were found in Antarctica and Argentinian Andes, respectively.

The importance of environmental factors for radioresistance was also confirmed by our results on the Antarctic endemic species *F. endolithicus* strains, living in the driest, coldest highly solar and UV radiation impacted environment and here resulted the far most radioresistant guild, together with *Cryomyces* strains living in the same desert (i.e. Antarctica). The relation occurring between the radioresistance and the adaptation to other environmental stressors may be mediated by both morphological and physiological properties of the fungal cells. Previous studies have indicated morphological characteristics of black fungi as possible determinants against environmental stressors (Kogej et al., 2006). Furthermore, the melanized cell wall was shown to play a key role in mediating the resistance of pigmented fungi to ionizing radiation (Dadachova et al., 2008; Shuryak et al., 2014). Recent advances showed various possible trends, represented by both linear and non-linear threshold correlations (Pacelli et al., 2018b; Schultzhaus et al., 2020). In our study, all the tested strains showed a non-linear exponential relationship between the gamma rays doses and the survival fraction, suggesting that radioresistance may be mediated by additional physiological and molecular processes differentially expressed in distinct species of black fungi (Robertson et al., 2012; Sharma et al., 2017; Romsdahl et al., 2020; Kreusch et al., 2021; Das et al., 2022; Kanekar & Kanekar, 2022). This may indicate how the response of microorganisms to ionizing radiation depends not only on the tested organisms, but also on the exposure conditions. In this perspective, future studies focusing on the genetic and molecular mechanisms mediating the radioresistance in black fungi should be performed. Our study confirms black fungi as excellent models for the study of the mechanisms used by eukaryotic cells to cope with ionizing radiation (Selbmann et al., 2018). We observed a general stunning radioresistance of most of the black fungi if compared to other microorganisms and some of them exceed the survival limits of the most radioresistant prokaryotic and eukaryotic forms of life (e.g. Cox et al., 2005; Daly, 2009; Sukhi et al., 2009; Singh et al., 2013; Krisko, 2013; Shuryak, 2019b). However, it should be highlighted that the response to ionizing radiation in microorganisms can vary markedly with the cellular state and exposure conditions (Hafer et al., 2010; Schultzhaus et al., 2020; Couceiro et al., 2021). Considering the dose rates recorded in the most radioactive environments on Earth, our results show how part of black fungi may hypothetically survive there even after thousands of years exposure to ionizing radiation in a silent desiccated state (Kashparov et al., 2018; Matsuo et al., 2019). Similarly, the radiation levels recorded in the interplanetary space and on the surface of celestial bodies that could host forms of life, such as Mars, are remarkably lower than those tested in our experiment (Zeitlin et al., 2013; Hassler et al., 2014; Inozemtsev et al., 2015).

Taken together, by simultaneously testing a broad set of ecologically and phylogenetically distinct fungal strains, the present study supplies the first characterization of the radioresistance in black fungi. Along with ecological information on the strains, these results allowed us to display how the radioresistance in black fungi may be the result of their adaptation to various environmental variables. From an evolutionary perspective, our findings demonstrate how the pressure exerted by a few stressors may induce mechanisms that enable the microorganisms to cope with a broader range of possible damaging factors. In the overall, this work may give new insights into the prediction of evolutionary responses to extreme environments and into the ability of life to adapt and persist in harsh terrestrial and extraterrestrial environments (e.g. Mars). This information represents the baseline for untangling the genomic and metabolomic traits underlying the radioresistance of black fungi, which may pave the way to untangle the limits of life in extreme radioactive environments.

## Acknowledgments

This research was financially supported by the Italian Space Agency (ASI), Life in Space – ASI N. 2019-3-U.0, and BioSigN MicroFossils – ASI N. 2018-6-U.0. We thank the Italian National Program of Antarctic Research (PNRA) and the Italian National Antarctic Museum “Felice Ippolito” (MNA) are acknowledged for funding the collection of Antarctic samples MNA-CCFEE and CCFEE. M.D-B. acknowledges support from the Spanish Ministry of Science and Innovation for the I+D+i project PID2020-115813RA-I00 funded by MCIN/AEI/10.13039/501100011033. M.D-B. is also supported by a project of the Fondo Europeo de Desarrollo Regional (FEDER) and the Consejería de Transformación Económica, Industria, Conocimiento y Universidades of the Junta de Andalucía (FEDER Andalucía 2014-2020 Objetivo temático “01 - Refuerzo de la investigación, el desarrollo tecnológico y la innovación”) associated with the research project P20_00879 (ANDABIOMA).

## Authors contributions

L.A., C.C., and L.S., designed the study and interpreted the results; L.A. performed molecular and statistical analysis and led the writing; all authors revised the manuscript and approved the final version.

## Data availability statement

All data generated or analyzed during this study are included in this published article and its supplementary information files.

## Conflict of interest

The authors declare no competing financial interests.

## Supporting information captions

**Figure S1**. Survival curves and calculated D10 values in strains of *Eurotomycetes*.

**Figure S2**. Survival curves and calculated D10 values in strains of lowly resistant *Dothideomycetes*

**Figure S3**. Survival curves and calculated D10 values in strains of highly resistant *Dothideomycetes*.

**Supplementary Table S1**. Fungal strains tested in the study and their response to radiation exposure in terms of D10 and the maximum survival dose.

**Supplementary Table S2**. Fungal strains and the localities from where they were collected and the related environmental metadata. AMT: Annual Mean Temperature; MTWAQ: Mean Temperature of the Warmest Quarter; MTCQ: Mean Temperature of the Coldest Quarter; AP: Annual Precipitation; PWEM: Precipitation of the Wettest Month; PDM: Precipitation of the Driest Month; PS: Precipitation Seasonality (Coefficient of Variation); PWEQ: Precipitation of the Wettest Quarter; PDQ: Precipitation of the Driest Quarter; PWAQ: Precipitation of the Warmest Quarter; PCQ: Precipitation of the Coldest Quarter; MDR: Mean Diurnal Range (Mean of monthly (maximum temperature - minimum temperature)); TS: Temperature Seasonality (standard deviation ×100); MTWAM: Maximum Temperature of the Warmest Month; MTCM: Minimum Temperature of Coldest Month; TAR: Temperature Annual Range (calculated as MTWAM - MTCM); MTWEQ: Mean Temperature of the Wettest Quarter; MTDQ: Mean Temperature of the Driest Quarter; IT: Isothermality (calculated as MDR/TAR) (×100); biome: biome type; country: country of isolation; KG climate: Köppen-Geiger climate classification subgroup.

## Notes

### Competing Interest Statement

The authors have declared no competing interest.

