## Supplementary figures and images for "Geography and environmental pressure are predictive of class-specific radioresistance in black fungi"

### Supplementary figure S2

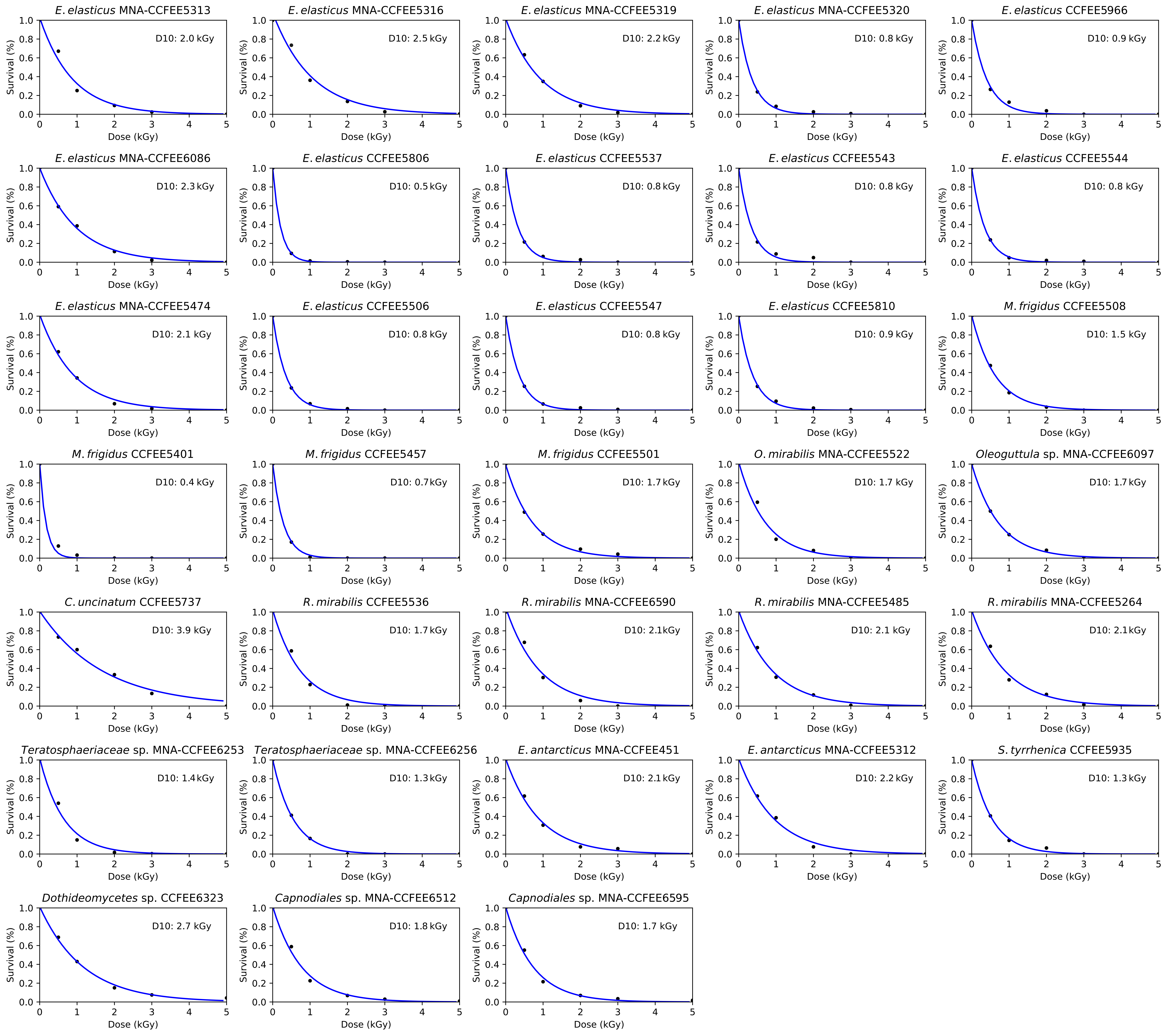

### Supplementary figure S3

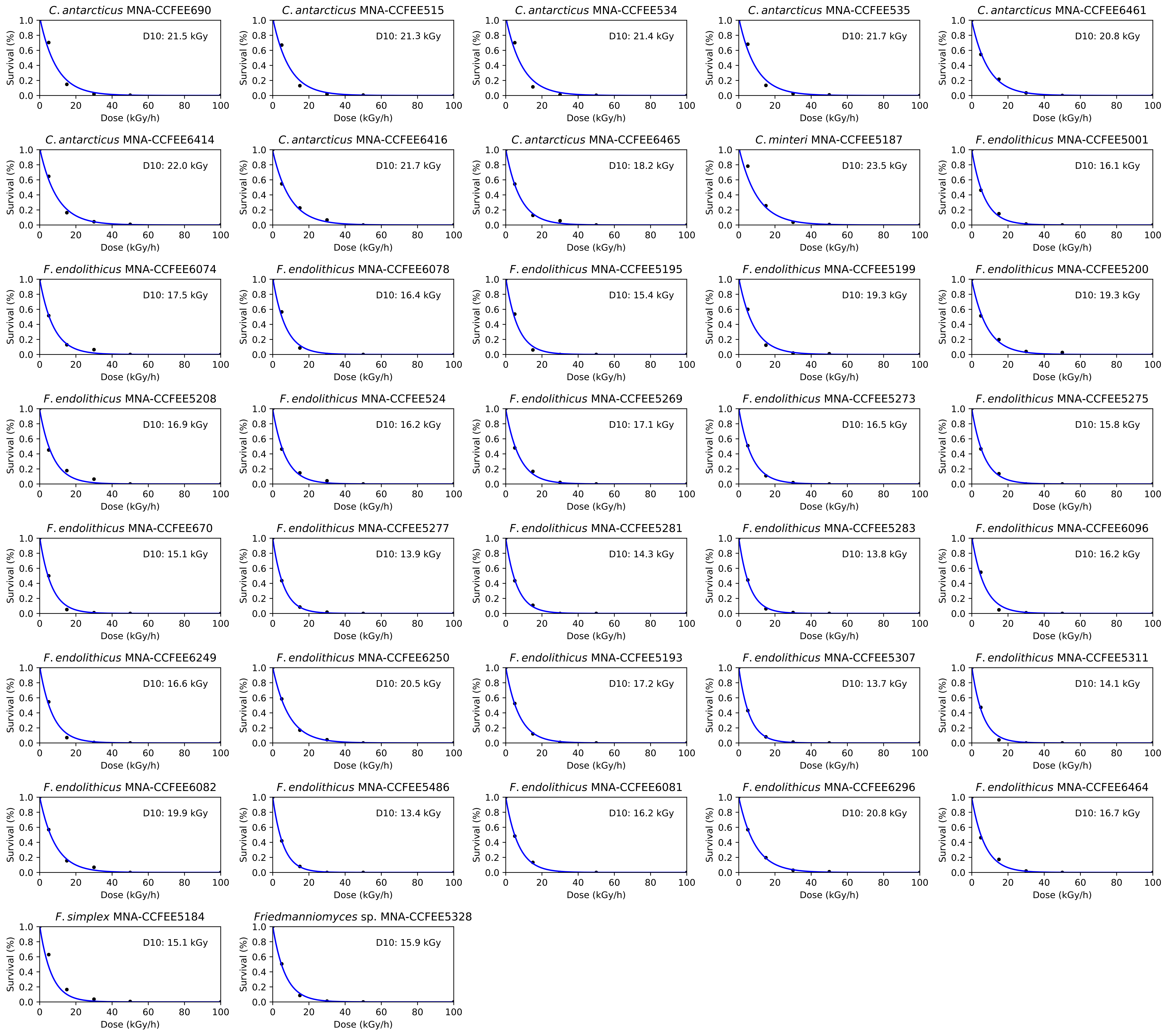
