## Supplementary Table S1 for "Geography and environmental pressure are predictive of class-specific radioresistance in black fungi"

| Class | Species | Strain | D10 (kGy) | Max dose (kGy) |
| --- | --- | --- | --- | --- |
| *Eurotiomycetes* | *Eurotiomycetes sp* | CCFEE6388 | 0.3 | 0.1 |
| *Eurotiomycetes* | *Chaetothyriales sp* | CCFEE6169 | 0.3 | 1 |
| *Eurotiomycetes* | *Exophiala xenobiotica* | CCFEE6043 | 0.4 | 1 |
| *Eurotiomycetes* | *Exophiala xenobiotica* | CCFEE5985 | 0.4 | 0.5 |
| *Eurotiomycetes* | *Exophiala xenobiotica* | CCFEE5816 | 0.4 | 2 |
| *Eurotiomycetes* | *Exophiala mesophila* | CCFEE6314 | 0.4 | 1 |
| *Eurotiomycetes* | *Exophiala xenobiotica* | CCFEE5877 | 0.4 | 3 |
| *Dothideomycetes* | *Meristemomyces frigidus* | CCFEE5401 | 0.4 | 1 |
| *Eurotiomycetes* | *Exophiala oligosperma* | CCFEE6327 | 0.4 | 1 |
| *Eurotiomycetes* | *Exophiala xenobiotica* | CCFEE6357 | 0.4 | 1 |
| *Eurotiomycetes* | *Exophiala xenobiotica* | CCFEE6059 | 0.5 | 1 |
| *Dothideomycetes* | *Elasticomyces elasticus* | CCFEE5806 | 0.5 | 2 |
| *Eurotiomycetes* | *Exophiala xenobiotica* | CCFEE5823 | 0.5 | 2 |
| *Eurotiomycetes* | *Exophiala xenobiotica* | CCFEE6068 | 0.5 | 1 |
| *Eurotiomycetes* | *Exophiala xenobiotica* | CCFEE6221 | 0.6 | 1 |
| *Eurotiomycetes* | *Exophiala xenobiotica* | CCFEE6233 | 0.7 | 1 |
| *Eurotiomycetes* | *Exophiala xenobiotica* | CCFEE5882 | 0.7 | 2 |
| *Dothideomycetes* | *Meristemomyces frigidus* | CCFEE5457 | 0.7 | 1 |
| *Eurotiomycetes* | *Exophiala xenobiotica* | CCFEE5874 | 0.7 | 1 |
| *Dothideomycetes* | *Meristemomyces frigidus* | CCFEE5508 | 0.7 | 2 |
| *Dothideomycetes* | *Elasticomyces elasticus* | CCFEE5537 | 0.8 | 2 |
| *Eurotiomycetes* | *Exophiala xenobiotica* | CCFEE6036 | 0.8 | 2 |
| *Eurotiomycetes* | *Exophiala bonariae* | CCFEE5792 | 0.8 | 1 |
| *Dothideomycetes* | *Elasticomyces elasticus* | CCFEE5543 | 0.8 | 2 |
| *Dothideomycetes* | *Elasticomyces elasticus* | CCFEE5544 | 0.8 | 3 |
| *Eurotiomycetes* | *Exophiala xenobiotica* | CCFEE5784 | 0.8 | 1 |
| *Dothideomycetes* | *Elasticomyces elasticus* | CCFEE5506 | 0.8 | 3 |
| *Eurotiomycetes* | *Exophiala xenobiotica* | CCFEE5811 | 0.8 | 1 |
| *Eurotiomycetes* | *Exophiala xenobiotica* | CCFEE6060 | 0.8 | 1 |
| *Dothideomycetes* | *Elasticomyces elasticus* | CCFEE5320 | 0.8 | 3 |
| *Dothideomycetes* | *Elasticomyces elasticus* | CCFEE5547 | 0.8 | 3 |
| *Eurotiomycetes* | *Exophiala xenobiotica* | CCFEE5801 | 0.9 | 1 |
| *Dothideomycetes* | *Elasticomyces elasticus* | CCFEE5810 | 0.9 | 3 |
| *Dothideomycetes* | *Elasticomyces elasticus* | CCFEE5966 | 0.9 | 2 |
| *Eurotiomycetes* | *Exophiala xenobiotica* | CCFEE6336 | 1 | 1 |
| *Eurotiomycetes* | *Exophiala xenobiotica* | CCFEE6142 | 1 | 2 |
| *Eurotiomycetes* | *Exophiala xenobiotica* | CCFEE6194 | 1 | 1 |
| *Eurotiomycetes* | *Exophiala xenobiotica* | CCFEE5819 | 1.1 | 1 |
| *Eurotiomycetes* | *Exophiala xenobiotica* | CCFEE6182 | 1.2 | 1 |
| *Dothideomycetes* | *Teratosphaeriaceae sp* | MNA-CCFEE6256 | 1.3 | 2 |
| *Dothideomycetes* | *Saxophila tyrrhenica* | CCFEE5935 | 1.3 | 2 |
| *Dothideomycetes* | *Recurvomyces mirabilis* | MNA-CCFEE6590 | 1.3 | 3 |
| *Dothideomycetes* | *Oleoguttula mirabilis* | MNA-CCFEE5522 | 1.3 | 3 |
| *Eurotiomycetes* | *Exophiala xenobiotica* | CCFEE6237 | 1.3 | 2 |
| *Eurotiomycetes* | *Exophiala xenobiotica* | CCFEE6196 | 1.3 | 1 |
| *Eurotiomycetes* | *Exophiala xenobiotica* | CCFEE6180 | 1.4 | 2 |
| *Dothideomycetes* | *Teratosphaeriaceae sp* | MNA-CCFEE6253 | 1.5 | 3 |
| *Dothideomycetes* | *Oleoguttula sp* | MNA-CCFEE6097 | 1.7 | 2 |
| *Dothideomycetes* | *Capnodiales sp* | MNA-CCFEE6595 | 1.7 | 5 |
| *Dothideomycetes* | *Meristemomyces frigidus* | CCFEE5501 | 1.7 | 5 |
| *Dothideomycetes* | *Recurvomyces mirabilis* | CCFEE5536 | 1.7 | 2 |
| *Dothideomycetes* | *Capnodiales sp* | MNA-CCFEE6512 | 1.8 | 5 |
| *Dothideomycetes* | *Elasticomyces elasticus* | MNA-CCFEE5313 | 2.0 | 3 |
| *Dothideomycetes* | *Extremus antarcticus* | MNA-CCFEE451 | 2.1 | 3 |
| *Dothideomycetes* | *Elasticomyces elasticus* | MNA-CCFEE5474 | 2.1 | 2 |
| *Dothideomycetes* | *Recurvomyces mirabilis* | MNA-CCFEE5485 | 2.1 | 5 |
| *Dothideomycetes* | *Elasticomyces elasticus* | MNA-CCFEE5319 | 2.2 | 3 |
| *Dothideomycetes* | *Extremus antarcticus* | MNA-CCFEE5312 | 2.2 | 2 |
| *Dothideomycetes* | *Elasticomyces elasticus* | MNA-CCFEE6086 | 2.3 | 3 |
| *Dothideomycetes* | *Elasticomyces elasticus* | MNA-CCFEE5316 | 2.5 | 5 |
| *Dothideomycetes* | *Dothideomyces sp* | CCFEE6323 | 2.7 | 5 |
| *Dothideomycetes* | *Recurvomyces mirabilis* | MNA-CCFEE5264 | 2.7 | 3 |
| *Dothideomycetes* | *Coniosporium uncinatum* | CCFEE5737 | 3.9 | 3 |
| *Dothideomycetes* | *Friedmanniomyces endolithicus* | CCFEE5486 | 13.4 | 15 |
| *Dothideomycetes* | *Friedmanniomyces endolithicus* | MNA-CCFEE5307 | 13.7 | 15 |
| *Dothideomycetes* | *Friedmanniomyces endolithicus* | MNA-CCFEE5283 | 13.8 | 30 |
| *Dothideomycetes* | *Friedmanniomyces endolithicus* | MNA-CCFEE5277 | 13.9 | 30 |
| *Dothideomycetes* | *Friedmanniomyces endolithicus* | MNA-CCFEE5311 | 14.1 | 15 |
| *Dothideomycetes* | *Friedmanniomyces endolithicus* | MNA-CCFEE5281 | 14.3 | 15 |
| *Dothideomycetes* | *Friedmanniomyces simplex* | MNA-CCFEE5184 | 15.1 | 15 |
| *Dothideomycetes* | *Friedmanniomyces endolithicus* | MNA-CCFEE670 | 15.1 | 30 |
| *Dothideomycetes* | *Friedmanniomyces endolithicus* | MNA-CCFEE5195 | 15.4 | 15 |
| *Dothideomycetes* | *Friedmanniomyces endolithicus* | MNA-CCFEE5275 | 15.8 | 15 |
| *Dothideomycetes* | *Friedmanniomyces sp.* | MNA-CCFEE5328 | 15.9 | 30 |
| *Dothideomycetes* | *Friedmanniomyces endolithicus* | MNA-CCFEE5001 | 16.1 | 30 |
| *Dothideomycetes* | *Friedmanniomyces endolithicus* | MNA-CCFEE6096 | 16.2 | 30 |
| *Dothideomycetes* | *Friedmanniomyces endolithicus* | MNA-CCFEE524 | 16.2 | 30 |
| *Dothideomycetes* | *Friedmanniomyces endolithicus* | MNA-CCFEE6081 | 16.2 | 15 |
| *Dothideomycetes* | *Friedmanniomyces endolithicus* | MNA-CCFEE6078 | 16.4 | 15 |
| *Dothideomycetes* | *Friedmanniomyces endolithicus* | MNA-CCFEE5273 | 16.5 | 30 |
| *Dothideomycetes* | *Friedmanniomyces endolithicus* | MNA-CCFEE6249 | 16.6 | 30 |
| *Dothideomycetes* | *Friedmanniomyces endolithicus* | MNA-CCFEE6464 | 16.7 | 30 |
| *Dothideomycetes* | *Friedmanniomyces endolithicus* | MNA-CCFEE5208 | 16.9 | 30 |
| *Dothideomycetes* | *Friedmanniomyces endolithicus* | MNA-CCFEE5269 | 17.1 | 30 |
| *Dothideomycetes* | *Friedmanniomyces endolithicus* | MNA-CCFEE5193 | 17.2 | 30 |
| *Dothideomycetes* | *Friedmanniomyces endolithicus* | MNA-CCFEE6074 | 17.5 | 30 |
| *Dothideomycetes* | *Cryomyces antarcticus* | MNA-CCFEE6465 | 18.2 | 30 |
| *Dothideomycetes* | *Friedmanniomyces endolithicus* | MNA-CCFEE5200 | 19.3 | 50 |
| *Dothideomycetes* | *Friedmanniomyces endolithicus* | MNA-CCFEE5199 | 19.3 | 50 |
| *Dothideomycetes* | *Friedmanniomyces endolithicus* | MNA-CCFEE6082 | 19.9 | 30 |
| *Dothideomycetes* | *Friedmanniomyces endolithicus* | MNA-CCFEE6250 | 20.5 | 50 |
| *Dothideomycetes* | *Friedmanniomyces endolithicus* | MNA-CCFEE6296 | 20.8 | 50 |
| *Dothideomycetes* | *Cryomyces antarcticus* | MNA-CCFEE6461 | 20.8 | 30 |
| *Dothideomycetes* | *Cryomyces antarcticus* | MNA-CCFEE515 | 21.3 | 50 |
| *Dothideomycetes* | *Cryomyces antarcticus* | MNA-CCFEE534 | 21.4 | 50 |
| *Dothideomycetes* | *Cryomyces antarcticus* | MNA-CCFEE690 | 21.5 | 50 |
| *Dothideomycetes* | *Cryomyces antarcticus* | MNA-CCFEE535 | 21.7 | 50 |
| *Dothideomycetes* | *Cryomyces antarcticus* | MNA-CCFEE6416 | 21.7 | 30 |
| *Dothideomycetes* | *Cryomyces antarcticus* | MNA-CCFEE6414 | 22.0 | 50 |
| *Dothideomycetes* | *Cryomyces minteri* | MNA-CCFEE5187 | 23.5 | 50 |

**Table 1.** Tested fungal strains listed by their D10 values. The maximum doses at which colony growth was observed is also shown (Max dose (kGy)).
